# Modeling the effect of temperature on species coexistence

**DOI:** 10.1101/2025.11.04.686381

**Authors:** Serguei Saavedra, Yuguang Yang, José Ignacio Arroyo, Chris P. Kempes

## Abstract

1. Temperature reshapes organismal performance and species interactions, yet competition models rarely treat the declining warm side of thermal performance curves explicitly, even though that regime is likely to become increasingly important under climate warming.
2. We develop a density-dependent generalized Lotka–Volterra framework for pairwise exploitative competition in which effective competition coefficients depend on relative temperature-dependent consumption, and we analyse coexistence using the structural feasibility domain.
3. Within this model class, one result is robust: pairwise structural coexistence is maximized when competitors have equal effective consumption, so with symmetric overlap the temperature of maximal coexistence is the temperature at which their consumption thermal performance curves intersect.
4. Stronger ecological predictions are parameter dependent. In particular, whether coexistence declines more steeply under warming than under cooling, and whether the coexistence maximum for species with different thermal optima is shifted toward the lower optimum, depend on thermal-curve shape, relative peak heights, baseline competition, resource overlap, and any temperature dependence in carrying capacities.
5. For the illustrative asymmetric unimodal curves analysed here, shared optima produce maximal coexistence near the optimum and a steeper decline under warming than under cooling, whereas different optima produce an intermediate coexistence peak biased toward the lower optimum. A qualitative comparison with classic *Drosophila* pairwise competition experiments at 19°C and 25°C is consistent with these expectations.
6. Because the empirical comparison relies on broad thermal categories, unequal species groups, and binary pairwise outcomes, it should be interpreted as a proof of concept rather than a definitive quantitative test. More generally, our results clarify which aspects of temperature-dependent pairwise coexistence are robust consequences of thermal asymmetry and which are contingent on parameterization.

## Introduction

Global warming is reshaping the distribution and abundance of species worldwide, driving shifts in composition and altering ecosystem properties (Parmesan and Yohe, 2003; Walther et al., 2002; Pecl et al., 2017). Many of these changes are mediated by species interactions, because warming modifies the rates at which organisms grow, consume resources, and suppress competitors (Tylianakis et al., 2008; Alexander et al., 2015; Tabi et al., 2020). Predicting those outcomes remains difficult because interaction strengths, demographic rates, and environmental supply can all respond to temperature, often nonlinearly. A useful starting point is therefore to isolate one tractable mechanism at a time and ask how it changes the conditions under which competitors can coexist.

Thermal performance curves (TPCs) provide the basic organismal link. Across many traits and taxa, performance typically rises with temperature up to an optimum and then declines sharply above it (Dell et al., 2011; Angilletta, 2009). Recent work suggests that much of this variation may collapse onto a parsimonious asymmetric form after appropriate rescaling (Arnoldi et al., 2025). Yet most theoretical studies of temperature-dependent competition emphasize the rising limb of the TPC or rely on monotonic thermal responses (Amarasekare and Savage, 2012; Sunday et al., 2019). That focus is understandable: many experiments are conducted below the thermal optimum, and in variable environments the temperature maximizing long-term performance can lie below the instantaneous optimum because of Jensen’s inequality (Martin and Huey, 2008). Nonetheless, the declining warm side of the TPC is ecologically central, especially for ectotherms already living close to their thermal optima (Deutsch et al., 2008; Vasseur et al., 2014). If climate warming increasingly pushes populations into that regime, supra-optimal temperatures may alter coexistence even when thermal effects near the optimum appear modest.

The idea that *thermal asymmetry* between interacting species can qualitatively change ecological dynamics is already well developed for consumer–resource systems (Dell et al., 2014; Vasseur and McCann, 2005). Competition should be equally sensitive to such asymmetry, because exploitative competitors differ in how temperature changes their rates of resource uptake and thus their effective suppressive effects on one another. However, the connection between full unimodal TPCs and competitive coexistence remains comparatively underdeveloped. In particular, it is not always clear what a temperature-dependent coexistence metric means biologically, whether pairwise results are robust to parameterization, and how pairwise results should and should not be extrapolated to richer communities.

Here we address that gap by focusing intentionally on *pairwise* competitive coexistence. We embed temperature-dependent resource consumption into a density-dependent generalized Lotka– Volterra (gLV) competition model and summarize coexistence with the structural feasibility domain (Saavedra et al., 2017, 2025). We do not seek a parameter-free universal forecast. Instead, we separate a robust geometric prediction from more contingent ecological patterns. The robust result is that, in the two-species case, coexistence is maximized when the competitors’ temperature-dependent consumptions are equal up to any fixed overlap asymmetry, so the co-existence peak occurs where the effective consumption ratio matches that asymmetry. We then use asymmetric unimodal TPCs to illustrate how stronger patterns can emerge under a biologically motivated parameter regime, quantify how often those stronger patterns persist in a broad numerical sensitivity analysis, and compare the resulting qualitative expectations with classic pairwise competition data from *Drosophila* (Gilpin et al., 1986). The empirical comparison is deliberately cautious: the data provide broad thermal categories rather than direct TPC measurements, the temperate and tropical groups are unequal in size, and the outcomes are binary pairwise end states. We therefore complement the main comparison with a size-matched tropical-species downsampling analysis. The contribution of the paper is a transparent theory of how thermal asymmetry can reshape pairwise competition beyond thermal optima, together with explicit tests of which conclusions are robust, which are parameter contingent, and which are sensitive to empirical group imbalance.

## Methods

### Temperature-dependent competition model

*We study a set of S* competitors with density-dependent gLV dynamics written in *r/K* form as

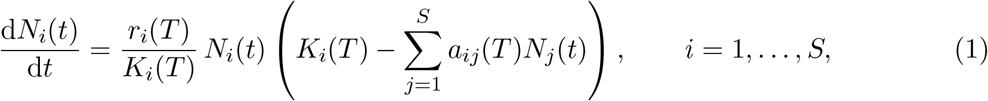

where *N*_*i*_(*t*) is the abundance or biomass of species *i, r*_*i*_(*T*) is its intrinsic growth rate, *K*_*i*_(*T*) is its carrying capacity, and *a*_*ij*_(*T*) is the per-capita effect of species *j* on species *i*, with *a*_*ii*_ = 1. Equation (1) is equivalent to the per-capita growth form

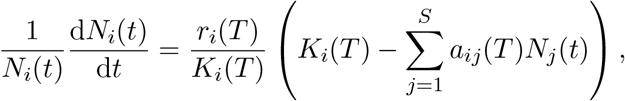

which makes explicit that density dependence enters through the focal population as well as through interspecific competition. Throughout, we interpret coexistence conditionally on *r*_*i*_(*T*) *>* 0 for all focal species over the temperatures considered. That is, the structural analysis below asks when competitors can coexist *given* that each species remains demographically viable in isolation at the focal temperature.

A positive equilibrium of Eq. (1) satisfies

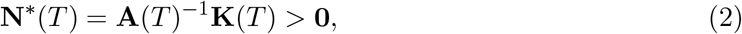

where **A**(*T*) is the interaction matrix and **K**(*T*) is the vector of carrying capacities. Temperature can therefore influence coexistence through multiple pathways: it can change demographic viability through *r*_*i*_(*T*), environmental supply through *K*_*i*_(*T*), and competitive asymmetry through **A**(*T*). Our focus is the last pathway. The coexistence metric below is used to isolate how temperature-dependent interaction asymmetry changes the set of relative environments compatible with coexistence; any particular temperature-dependent path of **K**(*T*) can then be interpreted relative to that set.

### Thermal asymmetry through resource consumption

To connect exploitative competition to temperature, we model interspecific competition as an effective consequence of shared resource use. Let *C*_*i*_(*T*) denote the mean per-capita rate at which species *i* consumes the relevant shared resource pool at temperature *T* . We write the effective competition coefficient as

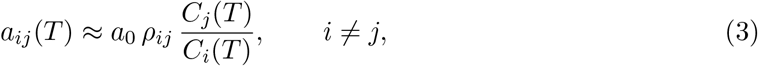

where *a*_0_ *>* 0 sets the baseline strength of interspecific competition and *ρ*_*ij*_ ∈ [0, 1] is a temperature-independent measure of resource-use overlap. Biologically, Eq. (3) says that, all else equal, a competitor that consumes more strongly from the shared resource pool imposes a larger suppressive effect on the other species.

Equation (3) should be interpreted as an *effective* reduction of a multi-resource exploitative competition model, not as a claim that competition for a single perfectly shared resource can by itself sustain coexistence. The role of *ρ*_*ij*_ is to absorb the temperature-independent structure of resource overlap and conversion efficiency; temperature then acts through the relative magnitude of total consumption. In the theoretical analysis below we set *ρ*_*ij*_ = 1 (or absorb it into *a*_0_) to isolate the consequences of thermal asymmetry. If temperature also changes diet composition, resource overlap, or resource dynamics, then additional temperature dependence would enter through *ρ*_*ij*_(*T*) and **K**(*T*); those extensions are important but are outside the scope of the present proof-of-concept model.

### Consumption thermal performance curves

The only requirement for the analytical results below is that each *C*_*i*_(*T*) be positive over the temperature range considered and vary unimodally with temperature. For numerical illustrations we therefore represent *C*_*i*_(*T*) with smooth asymmetric unimodal TPCs, parameterized by a lower thermal limit, an optimum temperature, an upper thermal limit, and a warm-side breadth parameter. This choice allows the common left-skewed shape emphasized in empirical work (Dell et al., 2011; Cruz-Loya et al., 2024; Arnoldi et al., 2025) while keeping the analysis interpretable in terms of optima, breadth, and relative asymmetry.

Crucially, the pairwise coexistence result depends only on the ratio *C*_*j*_(*T*)*/C*_*i*_(*T*), not on a particular mechanistic source of that ratio. Body size can therefore be incorporated when empirical information justifies it, for example by writing 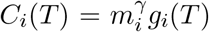 with an allometric exponent *γ* (Savage et al., 2004). *However, because the empirical Drosophila* comparison used here does not include body-size measurements, we do not make body-size-specific predictions in the main text. Body size is best viewed here as one optional determinant of the consumption curve rather than as a necessary ingredient of the theory.

### Parameter dependence and scope of inference

The numerical temperature–coexistence curves shown below are intended as *illustrations of a model regime*, not as claims of universal quantitative behavior. Within the present model class, Eq. (10) identifies a robust invariant: pairwise structural coexistence is maximized when the temperature-dependent consumptions are equal. By contrast, several stronger patterns are parameter contingent. These include the exact height of Ω, the steepness asymmetry between warming and cooling, the number and location of curve intersections, and whether the coexistence peak for species with different thermal optima lies closer to the lower or higher optimum. Those outcomes can vary with TPC skewness, relative peak heights, baseline competition strength *a*_0_, resource overlap *ρ*_*ij*_, relative body size when included, and any imposed temperature dependence in **K**(*T*).

We therefore interpret the theoretical figures as representative examples generated under a biologically motivated asymmetric unimodal parameterization rather than as exhaustive statements over parameter space. A full global sensitivity analysis would be the natural next step for quantitative forecasting, but the goal of the present manuscript is narrower: to identify the robust geometric mechanism and to distinguish it from the contingent predictions that depend on parameterization.

### Structural coexistence metric

For a fixed interaction matrix **A**, coexistence requires a positive equilibrium vector. Following structural coexistence theory (Saavedra et al., 2017, 2025), the set of carrying-capacity vectors that satisfy Eq. (2) defines the feasibility domain

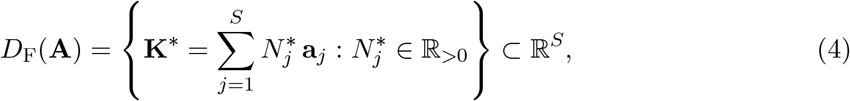

where **a**_*j*_ is column *j* of **A**. We quantify the size of this domain by the normalized measure

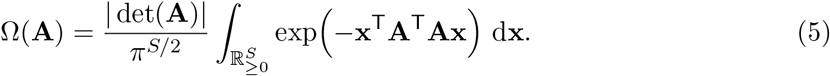

For fixed *S*, larger Ω means that coexistence is compatible with a broader range of *relative* carrying capacities. In other words, Ω is a structural robustness measure. It does *not* quantify equilibrium biomass magnitude, and in multispecies settings it does not count how many subsets of species coexist. Those distinctions matter when interpreting the metric beyond the pairwise case.

Because the empirical analysis here is pairwise, the two-species case is especially informative.

For

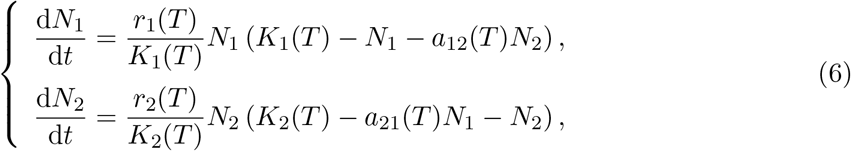

coexistence requires

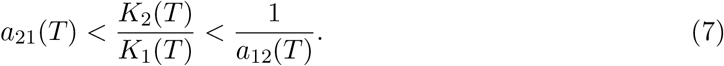

Thus, temperature-dependent carrying capacities do not invalidate the framework; rather, they define a temperature-dependent path for the ratio *K*_2_(*T*)*/K*_1_(*T*) through the feasible interval. The structural metric Ω(**A**) summarizes the width of that interval after averaging over all positive directions of **K**.

For two species, the normalized feasibility domain size can be written as

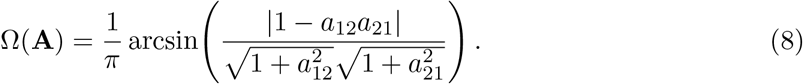

Now define the thermal asymmetry ratio

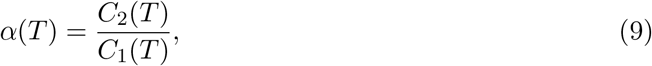

and substitute Eq. (3) with symmetric overlap into the pairwise interactions, so that *a*_12_(*T*) = *a*_0_*α*(*T*) and *a*_21_(*T*) = *a*_0_*/α*(*T*). Equation (8) is then maximized at *α*(*T*) = 1, implying

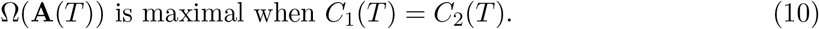

If the two consumption curves intersect at a unique temperature *T*_*×*_, then pairwise structural coexistence is maximal at *T*_*×*_. This provides the key interpretation used throughout the Results: temperature promotes coexistence when it reduces consumption asymmetry and suppresses coexistence when it amplifies that asymmetry.

This pairwise expression also yields a compact analytical sensitivity analysis. Let

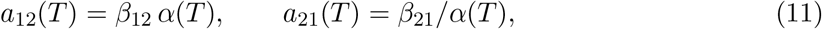

where *α*(*T*) = *C*_2_(*T*)*/C*_1_(*T*) and *β*_12_, *β*_21_ *>* 0 are temperature-independent multiplicative factors capturing baseline competition and fixed asymmetries in resource overlap. Substituting Eq. (11) into Eq. (8) shows that Ω is maximized at

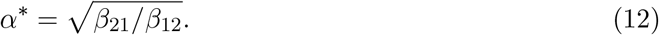

Thus, changing a common multiplicative factor such as *a*_0_ or symmetric temperature-independent overlap changes the *height* of Ω but not the temperature at which Ω is maximal. By contrast, fixed asymmetry in overlap shifts the peak away from *C*_1_(*T*) = *C*_2_(*T*) to the temperature at which 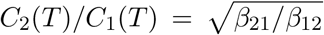. Temperature-dependent carrying capacities play a different role: they do not alter Ω(**A**(*T*)) itself, but they do change whether the realized path of *K*_2_(*T*)*/K*_1_(*T*) falls inside the feasible interval in Eq. (7). We use these analytical sensitivities to distinguish robust from contingent predictions below.

### Empirical comparison with *Drosophila*

To compare these predictions qualitatively with data, we analyzed the classic laboratory competition dataset of Gilpin et al. (1986). We restricted attention to the thick-food pairwise experiments run at 19°C and 25°C, because those treatments were evaluated for the majority of pairs and are the most comparable to the simple competition model considered here. The dataset contains 25 species in total: 10 classified in the original source as temperate-distribution species and 15 as tropical-distribution species. We treat these categories as coarse proxies for lower and higher thermal optima, respectively. Throughout the main text we therefore refer to the categories as *temperate* and *tropical* ; species IDs are retained only in the matrices for indexing the original experiments.

The empirical comparison is intentionally conservative. First, distribution category is only a proxy for thermal optimum, not a direct TPC measurement. Second, the temperate and tropical groups are unequal in size (10 versus 15 species). The three pair classes therefore contain different numbers of ordered pairwise outcomes: 90 temperate–temperate comparisons (10 × 9), 210 tropical–tropical comparisons (15 × 14), and 150 mixed temperate–tropical comparisons (10 × 15). To reduce the most direct consequence of that imbalance, we summarize coexistence primarily as the *proportion* of observed pairwise outcomes within each pair class and report the corresponding raw counts in the Results. Third, the outcomes are binary pairwise end states rather than fitted interaction coefficients. For each ordered pair and temperature, we recorded the long-term outcome reported in the source matrices. When the source marked replicate reversals or occasional coexistence with an asterisk, we retained both the modal outcome and a binary flag for “coexistence observed at least once”. The primary analysis uses the latter as a permissive proof-of-concept endpoint; the modal endpoint was used as a sensitivity check.

We then compared three classes of pairs: temperate–temperate, tropical–tropical, and mixed temperate–tropical. The theoretical expectations are straightforward. If competitors share similar thermal optima, coexistence should be more frequent near that shared optimum. If competitors differ in thermal optima, coexistence should be maximized near the temperature at which their consumption curves intersect, which is generally expected to fall between the optima. Because the comparisons are made within pair classes, unequal group sizes affect the precision of the estimated proportions more than the sign of the contrast itself. The empirical analysis is not used to estimate the parameters of the model; it is used only to ask whether the observed qualitative ordering matches a plausible parameter regime of the pairwise theory.

### Global sensitivity analysis, realized K(*T*) paths, and size-matched downsampling

To quantify parameter dependence more directly, we performed three auxiliary analyses. First, we generated broad ensembles of asymmetric unimodal consumption curves using a piecewise-Gaussian parameterization,

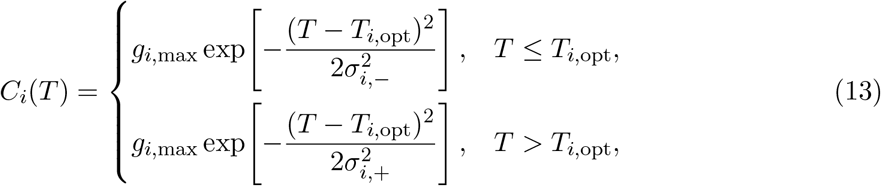

where *g*_*i*,max_ is the peak consumption rate and *σ*_*i*,−_ and *σ*_*i*,+_ are the cool-side and warm-side thermal breadths. This auxiliary parameterization was used only for sensitivity analysis; the analytical results above do not depend on it. For each draw we sampled *g*_*i*,max_ ∈ [0.7, 1.3], *σ*_*i*,−_ ∈ [3, 9], *σ*_*i*,+_ ∈ [1.5, 7], a common competition scale *β* ∈ [0.05, 0.95], and a fixed overlap asymmetry 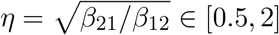. We then computed Ω(**A**(*T*)) on a dense temperature grid from 0 to 40°C. For 20,000 shared-optimum draws we recorded whether coexistence declined more steeply under warming than under cooling using equal 2°C perturbations around the maximizing temperature. For 20,000 different-optimum draws we recorded whether the maximizing temperature fell between the two optima and whether it was closer to the lower optimum. We also verified numerically, over 5,000 additional random draws, that the maximizing temperature matched the analytical condition in Eq. (12) to grid resolution.

Second, to study temperature-dependent carrying capacities without changing the structural metric itself, we sampled 5,000 linear trajectories for the log carrying-capacity ratio,

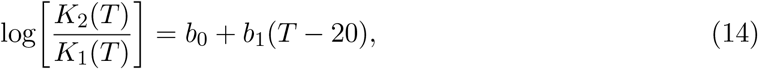

with *b*_0_ ∈ [−1, 1] and *b*_1_ ∈ [−0.12, 0.12]. For each path we evaluated the pairwise feasibility condition in Eq. (7) and computed the realized feasibility margin

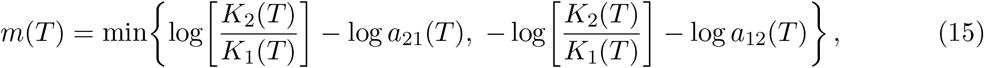

which is positive exactly when coexistence is realized. This separates the structural optimum of Ω(**A**(*T*)) from the temperature at which a particular environmental path actually maximizes realized feasibility.

Third, to evaluate whether the unequal numbers of temperate and tropical species could by themselves generate the empirical contrasts, we performed a size-matched downsampling analysis on the tropical species pool. Specifically, we digitized the tropical–tropical and mixed matrices plotted in Fig. 3 and enumerated all 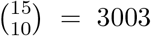 subsets of 10 tropical species. For each subset we recomputed the tropical–tropical coexistence proportion and the mixed temperate–tropical coexistence proportion at both temperatures. Because this downsampling uses the plotted matrices rather than the full raw transcription archive, we interpret it as a sign-robustness check rather than as a replacement for the main count summaries.

## Results

The pairwise analytical result in Eq. (10) provides the central organizing principle of the theory: coexistence is maximized when temperature equalizes the competitors’ consumptions. This equality rule is the robust prediction of the model class developed here. It does not depend on the detailed origin of the TPCs, only on how temperature changes the consumption ratio and therefore the competition asymmetry.

Figures 2a–b illustrate one biologically motivated parameter regime in which the two species share a common thermal optimum and have similar asymmetric unimodal TPCs. In that setting, the condition *C*_1_(*T*) = *C*_2_(*T*) is satisfied near the shared optimum, so the feasibility domain size peaks there. Because the illustrative TPCs are left-skewed, equal temperature departures above the optimum generate larger consumption asymmetries than equal departures below it. As a result, pairwise coexistence declines more steeply under warming than under cooling.

**Figure 1:**
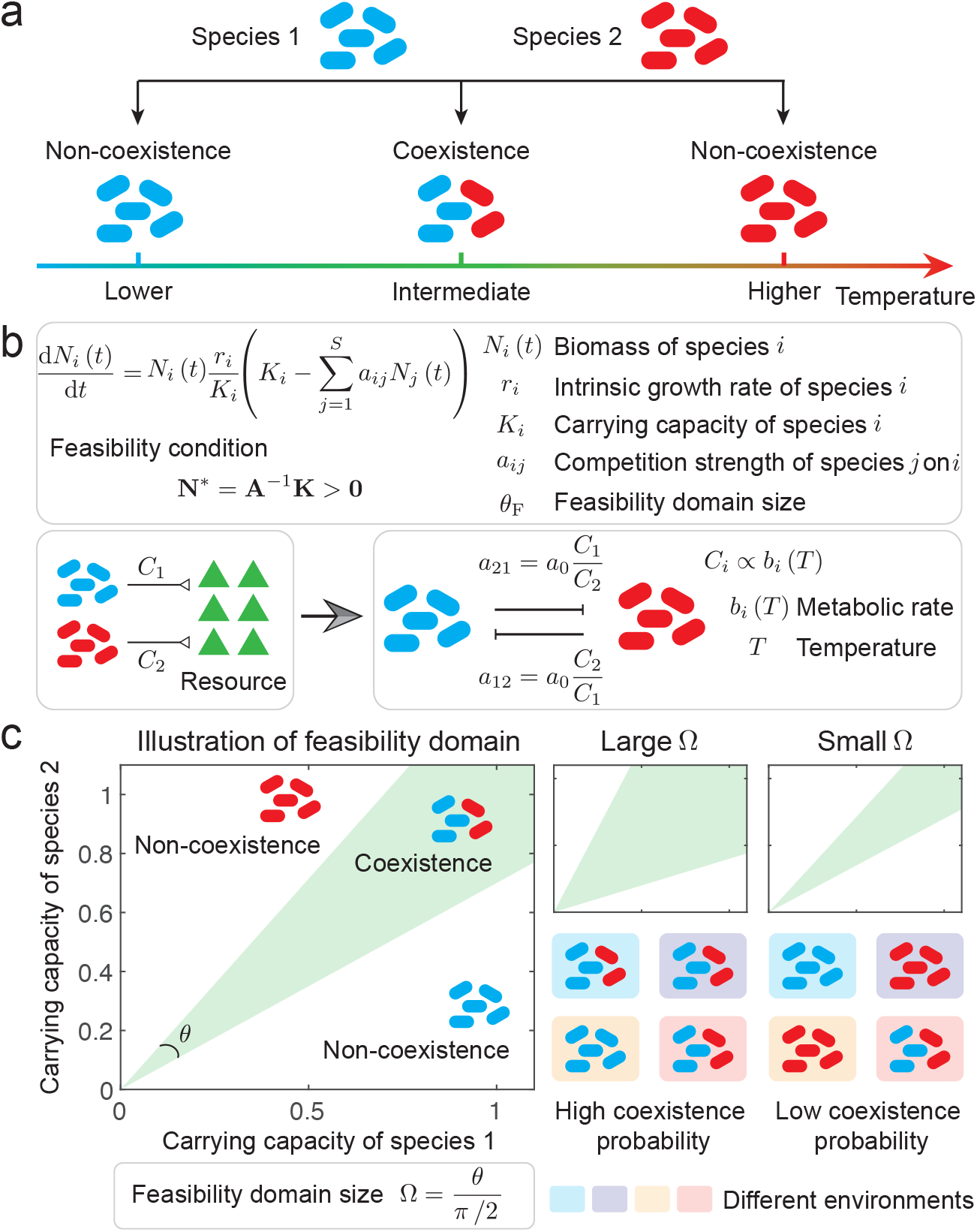
Temperature effects on pairwise competitive coexistence. **a**, Conceptual illustration of temperature-dependent coexistence for a pair of competitors with different thermal preferences. Coexistence occurs only over an intermediate temperature range. **b**, Temperature enters the density-dependent gLV competition model through temperature-dependent interaction coefficients derived from relative resource consumption. In the framework developed here, coexistence requires a positive equilibrium solution (**N**^*^ = **A**^−1^**K** *>* **0**). **c**, Structural coexistence theory. The feasibility domain is the set of relative carrying capacities that yield positive equilibria. Its normalized size (Ω) measures how robust coexistence is to variation in relative environmental supply: a larger Ω indicates that coexistence is compatible with a broader range of environments.

**Figure 2:**
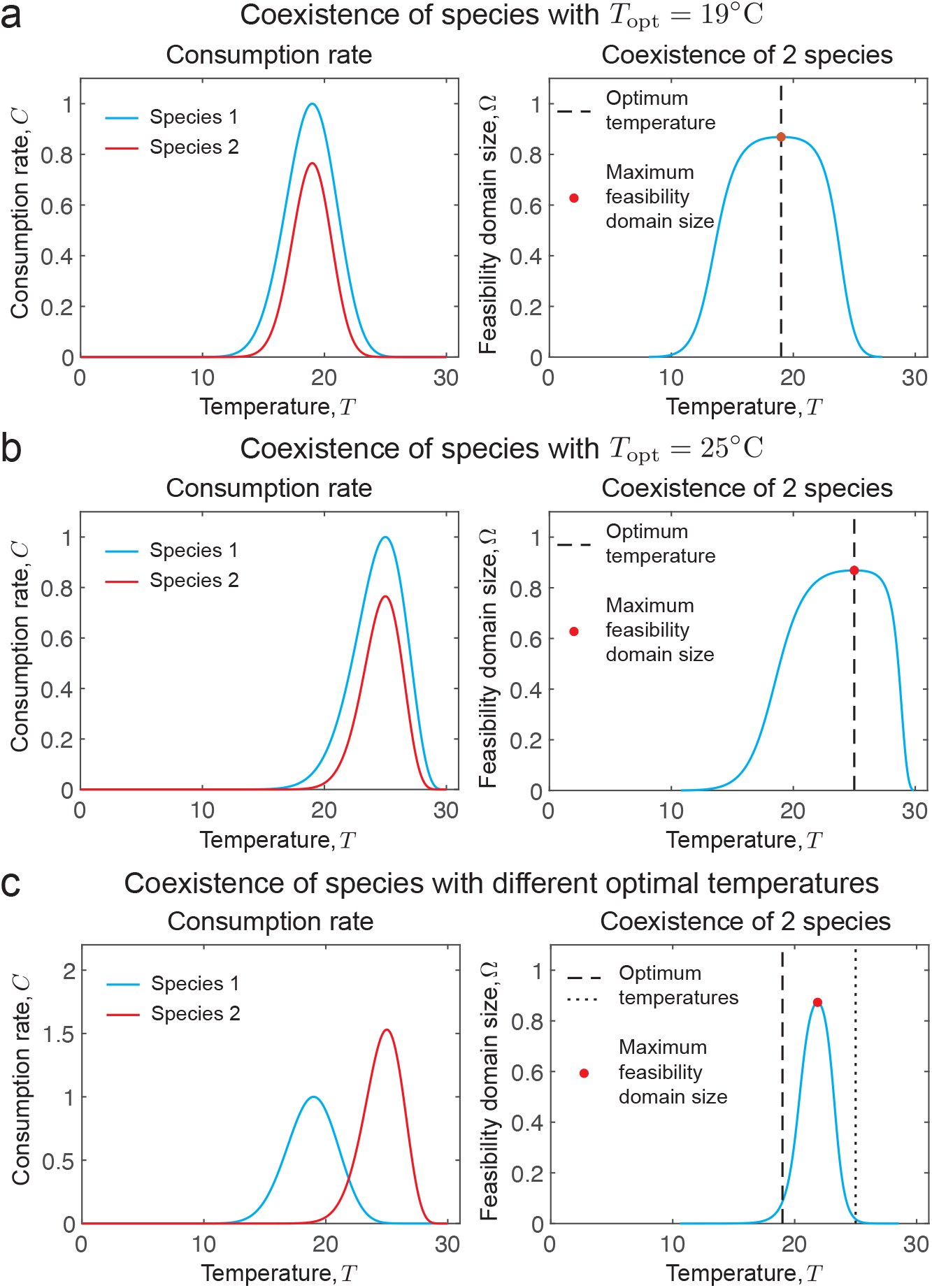
Illustrative parameter regimes for temperature-dependent pairwise coexistence. Panels **a** and **b** show species pairs with a shared thermal optimum; panel **c** shows a pair with different optima. In each panel, the left subplot gives the temperature-dependent consumption curves and the right subplot gives the corresponding feasibility-domain size. The robust pairwise prediction is that coexistence is maximal where the two consumption curves intersect, because that is where competitive asymmetry is minimized. The additional features shown here are parameter contingent. For the asymmetric unimodal curves in **a** and **b**, the intersection occurs near the shared optimum and coexistence declines more steeply under warming than under cooling. In **c**, the crossing occurs between the two optima and, for this illustrative parameterization, closer to the lower one. Different parameter values can alter the steepness and location of these peaks.

**Figure 3:**
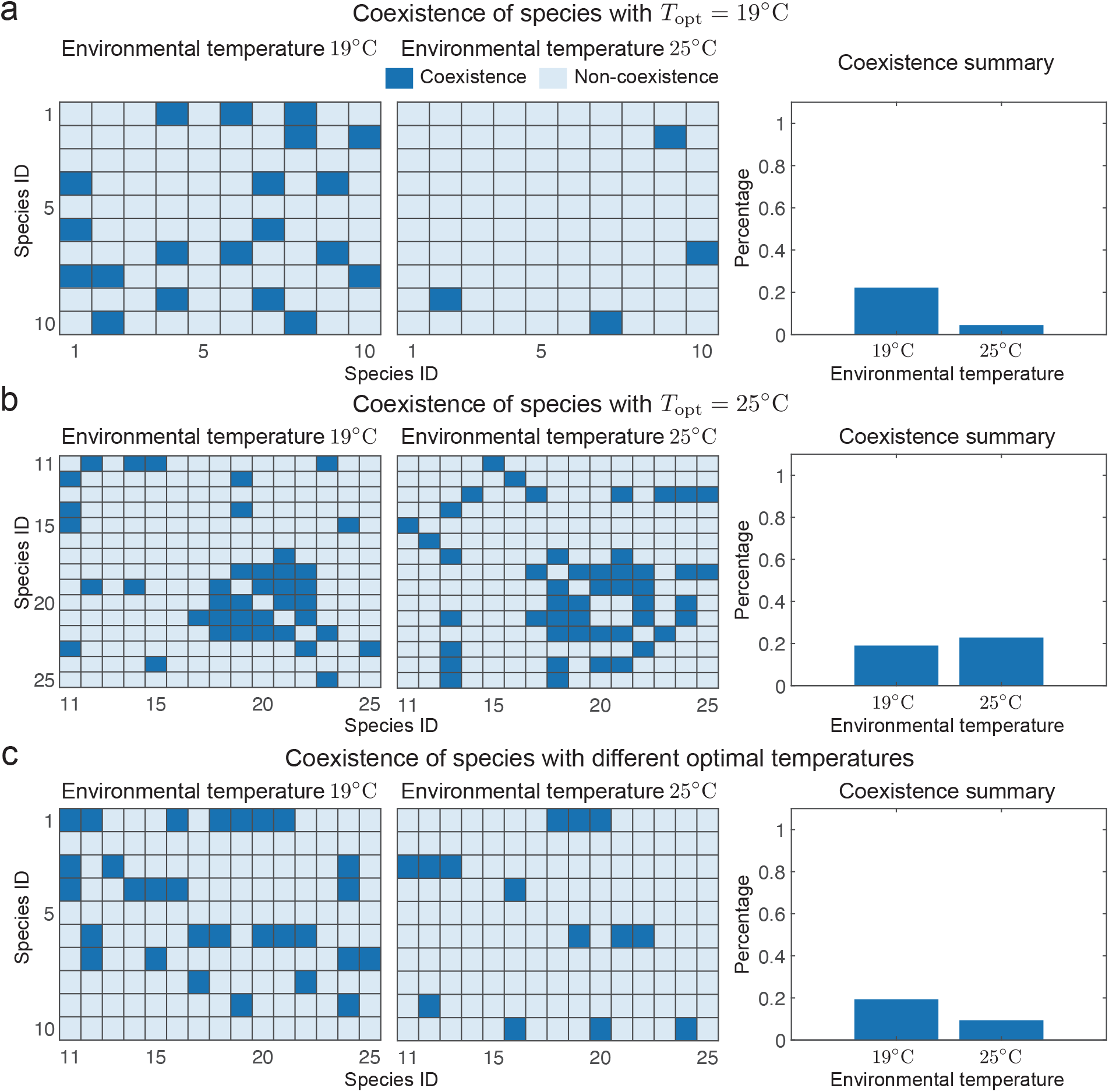
Qualitative comparison with pairwise *Drosophila* competition experiments. Panel **a** shows temperate–temperate pairs, panel **b** tropical–tropical pairs, and panel **c** mixed temperate–tropical pairs. The original dataset contains 10 temperate-distribution species and 15 tropical-distribution species. Matrix entries show pairwise coexistence outcomes at 19°C and 25°C; dark squares denote coexistence and light squares denote non-coexistence. The bars report *within-class proportions* of coexistence rather than raw counts, which reduces the most direct effect of the unequal group sizes; the corresponding counts are reported in Supplementary Table 2. The comparison is qualitative because the thermal categories are broad proxies for thermal optima and the experiments were conducted at only two temperatures, but the observed ordering is consistent with one plausible parameter regime of the pairwise theory.

When the competitors differ in thermal optima, the coexistence peak shifts away from either optimum and toward the temperature at which the two consumption curves cross. Figure 2c shows this case explicitly. For the illustrative parameterization used there, the crossing temperature lies between the two optima and closer to the lower one, so coexistence is highest at an intermediate but relatively cool temperature. This lower-optimum bias is therefore not a separate assumption; it emerges from the location of the TPC crossing for the chosen asymmetric curves.

These stronger directional patterns are not universal. Within the present framework, the exact shape of the temperature–coexistence curve depends on TPC skewness, relative peak heights, baseline competition strength, resource overlap, and any temperature dependence imposed on relative carrying capacities. Different parameterizations can flatten the warm–cool asymmetry, shift the location of the coexistence peak, or generate more than one crossing of the consumption curves. Figure 2 should therefore be read as an illustration of one plausible regime rather than as an exhaustive map of outcomes over parameter space.

The analytical sensitivity result in Eq. (12) helps organize these dependencies. Parameters such as a common baseline competition factor *a*_0_ or symmetric, temperature-independent overlap scale the magnitude of Ω without changing the temperature that maximizes it. By contrast, fixed asymmetries in overlap shift the maximizing temperature, and temperature-dependent carrying capacities can move a realized system into or out of the feasible interval even when Ω(**A**(*T*)) is unchanged. The robust prediction is therefore about how coexistence tracks effective interaction symmetry, whereas the directional warm-versus-cool asymmetry and the exact peak location remain contingent on parameterization.

The auxiliary numerical sensitivity analysis makes that contingency explicit (Supplementary Table 1). Across 5,000 random parameter draws, the temperature maximizing Ω matched the analytical condition 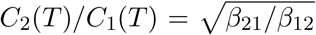 to grid resolution, with a median absolute difference of 0.00°C and a maximum discrepancy of 0.01°C. Beyond that invariant, however, the stronger directional patterns in Fig. 2 were far from universal. In 20,000 shared-optimum draws, coexistence declined more steeply under warming than under cooling in 54.1% of parameterizations overall, rising to 58.9% when both competitors had broader cool-side than warm-side thermal breadths and falling to 48.0% otherwise. In 20,000 different-optimum draws, the coexistence peak fell between the two optima in 58.2% of cases and was closer to the lower optimum in 61.3% of cases overall; when both competitors were left-skewed, the lower-optimum bias increased to 67.3%, whereas it was 53.2% otherwise. Figure 2 should therefore be interpreted as a representative regime rather than as a universal prediction.

Allowing **K**(*T*) to vary changed realized coexistence more strongly than structural coexistence. Under the sampled linear trajectories in Eq. (14), the structural optimum defined by Ω(**A**(*T*)) remained unchanged, but it fell inside the realized feasible region in only 59.3% of cases. Among the 4,805 sampled paths with at least one feasible temperature, temperature-dependent carrying capacities shifted the temperature maximizing the realized feasibility margin by a median of 2.28°C. This numerical check reinforces the conceptual distinction in Eq. (7): Ω(**A**(*T*)) quantifies the width of feasible relative environments, whereas **K**(*T*) determines whether a particular environmental trajectory actually traverses that region.

The *Drosophila* pairwise data show the same qualitative ordering as the illustrative regime. Using the permissive “coexistence observed at least once” endpoint, the within-class coexistence proportions correspond to 20/90 temperate–temperate ordered outcomes at 19°C versus 4/90 at 25°C, 41/210 tropical–tropical outcomes at 19°C versus 47/210 at 25°C, and 30/150 mixed outcomes at 19°C versus 13/150 at 25°C (Supplementary Table 2; Fig. 3). Thus, temperate– temperate pairs coexist more often at 19°C than at 25°C, tropical–tropical pairs show slightly higher coexistence at 25°C than at 19°C, and mixed temperate–tropical pairs coexist more often at 19°C than at 25°C. Because these summaries are reported as within-class proportions, they are not driven mechanically by the unequal number of temperate and tropical species, although the imbalance remains a limitation of the dataset. Taken together, the empirical comparison is consistent with the pairwise theory under a parameter regime in which coexistence is maximized near shared optima and, for mixed thermal types, at the cooler of the two experimental temperatures.

The empirical support should nevertheless be interpreted cautiously. The *Drosophila* analysis uses broad thermal categories rather than measured TPCs, the experimental temperatures are limited to two points, and the endpoint is a binary coexistence outcome. We therefore view Fig. 3 as a qualitative proof of concept rather than a quantitative parameter fit. Its value is that the direction of the observed contrasts matches the direction predicted by a plausible pairwise parameterization.

The size-matched downsampling analysis shows that the unequal sizes of the temperate and tropical pools do not fully explain the empirical ordering (Supplementary Table 3). Across all 3,003 subsets of 10 tropical species, mixed temperate–tropical pairs coexisted more often at 19°C than at 25°C. The tropical–tropical contrast was less stable but still usually preserved: coexistence at 25°C was at least as frequent as coexistence at 19°C in 76.1% of size-matched subsets and strictly higher in 66.1% of them. Unequal group sizes therefore remain a limitation, especially for the tropical–tropical contrast, but they do not mechanically generate the mixed-pair result.

## Discussion

This revision reframes the manuscript around a narrower but more defensible claim: asymmetric unimodal thermal responses can reshape *pairwise* competitive coexistence by changing the relative consumption rates that determine effective interaction strengths. Within that framing, the key robust result is simple and transparent. In a two-species exploitative competition model with fixed resource overlap, structural coexistence is maximized when the competitors consume equally, so the coexistence peak occurs at the temperature where their consumption TPCs intersect.

The broader ecological patterns are more conditional. Shared optima can favor coexistence near the optimum, different optima can shift the coexistence peak to an intermediate temperature, and left-skewed TPCs can make warming more disruptive than cooling. But these are regime-level outcomes, not universal laws. They depend on how temperature shapes the consumption curves, on the magnitude of baseline competition, on resource overlap, and on the temperature dependence of carrying capacities. The global sensitivity analysis makes this point concrete: the equality rule remained exact across the full parameter sweep, but the warm-side asymmetry and lower-optimum bias appeared only in majorities, not all, of the sampled systems. Stating the results this way makes the paper both more accurate and more useful: the theory identifies the invariant mechanism, while the numerical illustrations and sensitivity analysis show how particular parameter combinations translate that mechanism into observable patterns.

A second contribution is conceptual. The feasibility measure Ω is often described abstractly as the size of a feasibility domain; here its biological meaning is made explicit for pairwise competition. In the two-species case, Ω is the fraction of relative carrying-capacity ratios compatible with a positive equilibrium. Large Ω therefore means that coexistence is robust to uncertainty or variation in relative environmental supply. It does not mean that the equilibrium has larger biomass, nor does it measure how many subsets of species coexist in a larger assemblage. This distinction is especially important beyond the pairwise case and is one reason we avoid over-interpreting the present results as direct predictions for full community assembly.

The model also clarifies the role of temperature-dependent demography. Reviewers correctly noted that intrinsic growth rates and carrying capacities often vary strongly with temperature (Savage et al., 2004; Bernhardt et al., 2018). Our framework does not deny that point. Instead, it separates three questions. First, are species individually viable at the focal temperature? That question concerns *r*_*i*_(*T*) and is treated here as a condition for applying the coexistence analysis. Second, does the realized environment fall inside the feasible region? That question concerns the path of **K**(*T*). Third, how large is the set of relative environments that would permit coexistence? That is the interaction-centered question summarized by Ω(**A**(*T*)). By focusing on the third question, we isolate the effect of thermal asymmetry in competition coefficients while remaining explicit about what the framework does and does not capture. The sampled **K**(*T*) paths show why that distinction matters: the structural optimum remained unchanged, yet realized coexistence often shifted by several degrees and failed to include the structural optimum in roughly 40% of the sampled cases. A fuller resource-explicit model would allow resource growth, carrying capacities, and resource overlap to vary with temperature simultaneously.

The *Drosophila* comparison is best read in that light. It uses a valuable historical dataset, but it is limited by coarse thermal grouping, unequal numbers of temperate and tropical species, only two experimental temperatures, and binary pairwise outcomes. Those features are sufficient for a qualitative check of direction, but not for a strong quantitative test or for adjudicating among parameterizations. The size-matched downsampling analysis strengthens the mixed-pair contrast and shows that the tropical–tropical contrast is usually, but not invariably, preserved under equalized group sizes. Future empirical work should therefore measure competitor-specific TPCs directly, estimate how temperature changes both resource uptake and demographic rates, and test the theory across continuous temperature gradients. On the theory side, the present global sensitivity analysis is intentionally broad rather than exhaustive; a next step is to repeat the exercise in resource-explicit models with temperature-dependent overlap and carrying-capacity trajectories. Data that include body size would also allow a more direct evaluation of when size helps explain thermal asymmetry in competition and when it does not.

Finally, although the feasibility formalism extends naturally to multispecies systems, the present paper should be read primarily as a pairwise theory plus a pairwise empirical comparison. That narrower scope is a strength rather than a weakness. It makes the biological meaning of the metric transparent, avoids unsupported extrapolation from pairwise data to whole communities, and provides a clean baseline for future resource-explicit and multispecies extensions. In that sense, the main message is straightforward: to understand how warming changes competitive coexistence, one must pay attention not only to thermal optima, but also to the full asymmetric shape of the thermal performance curve beyond the optimum—and to which conclusions are robust versus parameter contingent.

## Acknowledgments

SS acknowledges support from the National Science Foundation under Grant No. DEB-2436069. JIA acknowledges support from the National Science Foundation under Grant No. 2133863, and from the Center for Mathematical Modeling (CMM), Grant FB210005, BASAL funds for Centers of Excellence from ANID-Chile.

## Data and code availability

No new data were generated in this paper. The *Drosophila* data were transcribed from Gilpin et al. (1986). Code to generate the numerical illustrations, compute feasibility measures, and reproduce the figures is available at https://github.com/MITEcology/Temperature_Coexistence.

## Ethics statement

This study involved no new experiments on animals or humans. All analyses were based on previously published laboratory data.

## Competing interests

The authors declare no competing interests.

## Supplementary Tables

The following tables are provided as supplementary material to support the sensitivity analyses and the empirical robustness checks discussed in the main text.

**Table 1:**
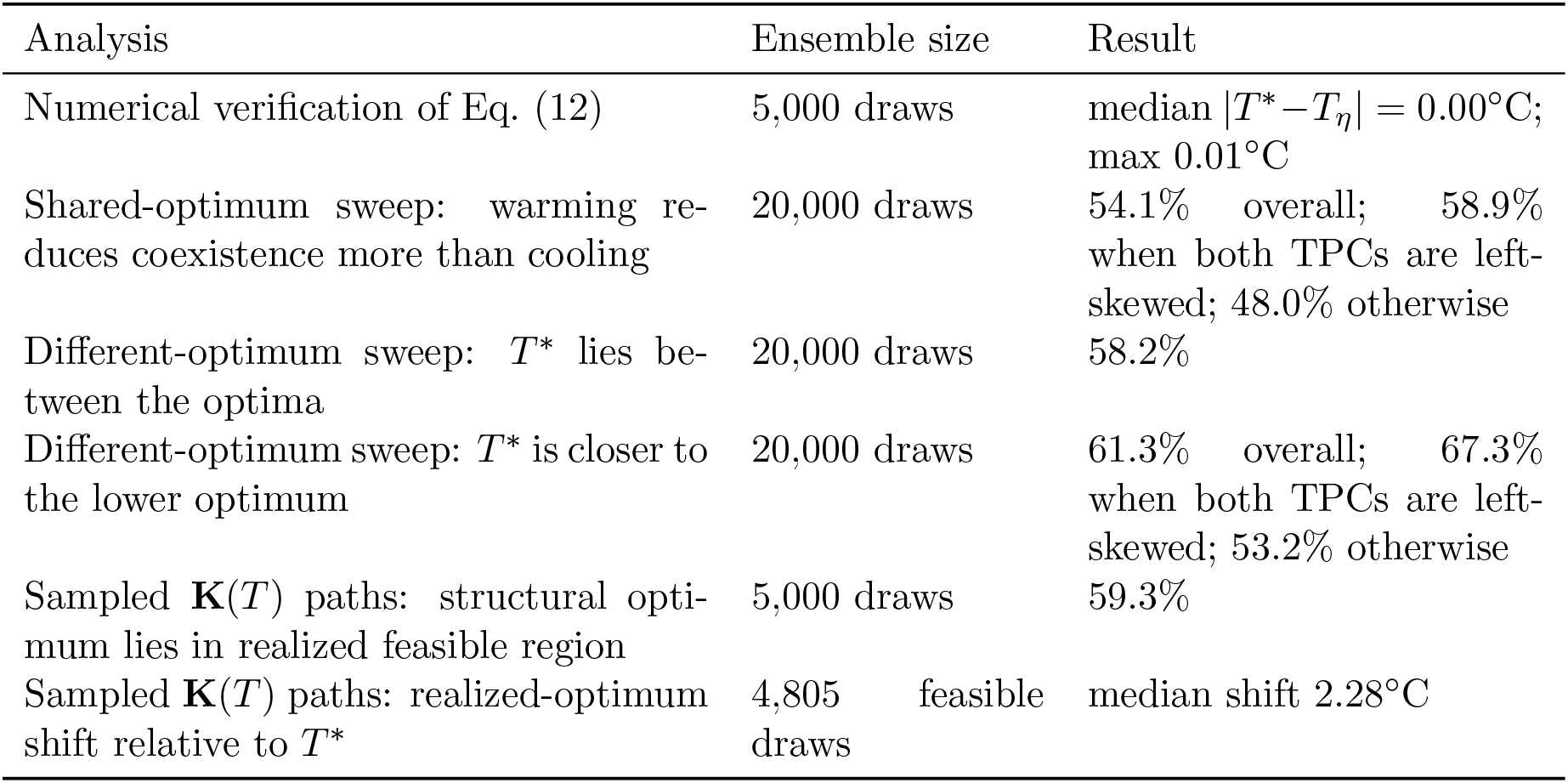
Summary of the auxiliary numerical sensitivity analyses. The global sweep used broad asymmetric piecewise-Gaussian consumption curves (Eq. (13)) and the sampled **K**(*T*) paths used Eq. (14).

**Table 2:** Observed coexistence counts in the *Drosophila* pairwise comparison, using the “coexistence observed at least once” endpoint. Denominators are the numbers of ordered pairwise outcomes within each class.

| Pair class | 19°C | 25°C |
| --- | --- | --- |
| Temperate–temperate | 20/90 | 4/90 |
| Tropical–tropical | 41/210 | 47/210 |
| Mixed temperate–tropical | 30/150 | 13/150 |

**Table 3:**
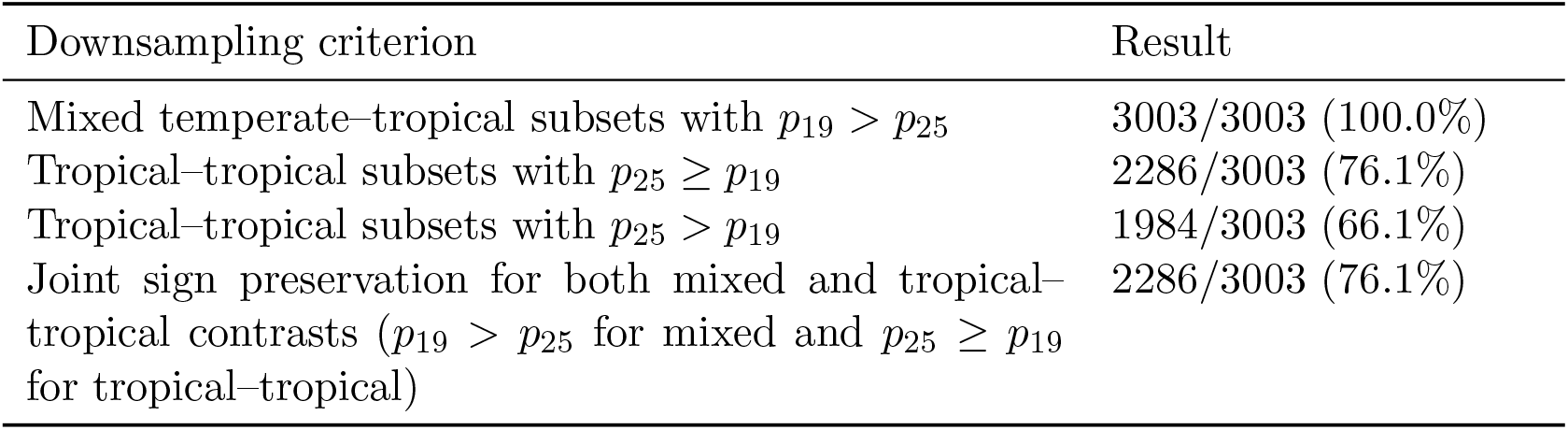
Size-matched downsampling of the tropical species pool using the digitized matrices plotted in Fig. 3. Each row summarizes the sign of the temperature contrast over all 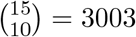 subsets of 10 tropical species.

## Notes

https://github.com/MITEcology/Temperature_Coexistence

### Competing Interest Statement

The authors have declared no competing interest.

### Summary of Updates

Corrected typos, added sensitivity analysis, expanded text

